# *jouvence*, a new human H/ACA snoRNA involves in the control of cell proliferation and differentiation

**DOI:** 10.1101/2020.06.17.157321

**Authors:** Flaria El-Khoury, Jérôme Bignon, Jean-René Martin

## Abstract

Small nucleolar RNAs (snoRNAs) are non-coding RNAs conserved from archeobacteria to mammals. In humans, various snoRNAs have been associated with pathologies as well as with cancer. Recently in *Drosophila*, a new snoRNA named *jouvence* has been involved in lifespan. Since snoRNAs are well conserved through evolution, both structurally and functionally, *jouvence* orthologue has been identified in human, allowing hypothesizing that *jouvence* could display a similar function (increasing healthy lifespan) in human. Here, we report the characterization of the human snoRNA-*jouvence*, which was not yet annotated in the genome. We show, both in stably cancerous cell lines and in primary cells, that its overexpression stimulates the cell proliferation. In contrast, its knockdown, by siRNA leads to an opposite phenotype, a decrease in cell proliferation. Transcriptomic analysis reveals that overexpression of *jouvence* leads to a dedifferentiation signature of the cells, a cellular effect comparable to rejuvenation. Inversely, the knockdown of *jouvence* leads to a decrease of genes involved in ribosomes biogenesis and spliceosome in agreement with the canonical role of a H/ACA box snoRNA. In this context, *jouvence* could represent a now tool to fight against the deleterious effect of aging, as well as a new target in cancer therapy.

## Introduction

Small nucleolar RNAs (snoRNAs), which consist of 60-300 nucleotides and conserved from archeobacteria to mammals, are non-coding RNAs localized in the nucleolus where they are associated with proteins to form small nucleolar RiboNucleoProtein (snoRNPs) [1,2]. They can be subdivided, based on conserved secondary structure and functional RNA motifs, in two major classes, the box C/D and the box H/ACA. They are predicted to guide nucleotide modifications mainly of ribosomal RNAs (rRNAs). Most box C/D performs the 2’-O-methylation of RNAs through fibrillarin, while the box H/ACA are known to convert the uridine to pseudouridine through dyskerin [1-4]. However, as more recently reported, H/ACA snoRNA seems to also pseudouridinylate other RNA substrates, such as mRNA and long-non-coding-RNAs (lncRNAs) [5]. In addition, few other functions have also been assigned to H/ACA box snoRNAs, as for instance chromatin remodelling [5]. In humans, some pathologies have been associated with various H/ACA snoRNAs, as the congenital dyskeratose, in which patients have shorter telomeres [6,7], while the snoRNA HBII-52, a human C/D box, has been involved in the alternative splicing of the serotonine receptor 2C [8,9].

In addition to their reported role in some specific pathologies, recent reports suggest that snoRNAs have also tumor-suppressive or oncogenic functions in various cancer types [10]. Indeed, several snoRNAs have been reported to participate in many biological cancer processes, as inactivation of growth suppressors and cell death, activation of invasion and metastasis, as well as sustained proliferative signalling [11]. Moreover, some evidences suggest that snoRNA could also play a role in cancer stem cells (CSC), as these last have the capacities of self-renewal, differentiation and tumorigenicity [12]. Therefore, in this context, snoRNAs could have potential applications for cancer diagnosis and therapy.

In the last few years, in *Drosophila*, we have identified a new H/ACA box small nucleolar RNA (snoRNA) (a non-coding RNA), named *jouvence* (*jou*), the deletion of which reduces lifespan [13]. Inversely, its overexpression increases lifespan. In *Drosophila, jouvence* is required in the epithelium of the gut, and more precisely in enterocytes, while its deletion yields to the gut hyperplasia/dysplasia in aged flies [13]. Since the snoRNAs are well conserved throughout evolution, both structurally and functionally [1-4], we have identified its orthologues in mouse and human. The mouse genome contains two snoRNAs *jouvence* genes, whereas only one copy has been identified in human, located on chromosome 11. RT-PCR show that, both in mouse and human, *jouvence* orthologues are expressed suggesting that they might be functional [13]. Thus, these results allow to hypothesize that *jouvence* could display a similar function (increasing healthy lifespan) in mammals, including human.

Here, we have characterized the newly identified snoRNA-*jouvence* in human, which was not yet annotated. First, we have defined its genomic map as well as its predicted secondary structure. Second, we have shown, by RT-qPCR, that several cell types expressed *jouvence*. We have also shown that its overexpression importantly stimulates the proliferation of the cells, both in immortalized cancerous cells lines, as HCT116 and Caco-2, as well as on non-cancerous cell lines as HEK293 and RPE1, and even in primary cells as HUVEC. Inversely, its knockdown, by transitory transfection with siRNA leads to an opposite phenotype, inhibiting the proliferation of the cells. In a step further, a transcriptomic analysis (RNA-Seq) performed on HCT116 overexpressing cells reveals a signature of dedifferentiation of the cells, compatible with a rejuvenation. Finally, a similar transcriptomic analysis performed on the cells in which *jouvence* is knocked-down by siRNA reveals a typical and canonical snoRNA phenotype, meaning a strong decrease of the ribosome biogenesis as well as a decrease of the spliceosome pathways. Therefore, since *jouvence* depletion reduces cancer cell proliferation, we hypothesis that it could represent a good candidate to fight against cancer.

## Results

### Genomic map, structure, and localization of the human snoRNA-*jouvence*

Based on the tertiary (3D) structure of the *Drosophila* snoRNA-*jouvence* through the uses of the infernal software, we have identified the orthologue of *jouvence* in mouse and human, which was not yet annotated (13]. Human *jouvence* (shortly named *h-jou*) is located on the chromosome 11 (GRCh38.p12) in a long intron of the TEA domain family member 1 gene (named TEAD1) (Figure 1A). The human genome contains a single copy of *jouvence*, while two copies have been identified in the mouse genome (for the human sequence of *jouvence*, see Figure 10 in Soulé et al., 2020) [13]. Since *jouvence* possesses a primary sequence corresponding to a H/ACA box, its secondary structure has been deduced from the UNAFold software, and as expected is folded to form a double hairpin (Figure 1B), a typical structure of the H/ACA box snoRNA [1,2]. In a step further, the expression level of *jouvence* was determined by RT-qPCR, in some well-established human cell lines (Figure 1C). The snoRNA is weakly detected in the nine tested cell lines (first Ct around 30 to 33). Expression levels were close for all the cell lines, with HEK293 cells expressing a little bit more *jouvence* compared to the other cell lines, with a dCt (Ct snoRNA - Ct reference gene GAPDH) around 10. The U87-MG glioblastoma cells had the lowest expression level with a dCt around 15.

**Figure 1).**
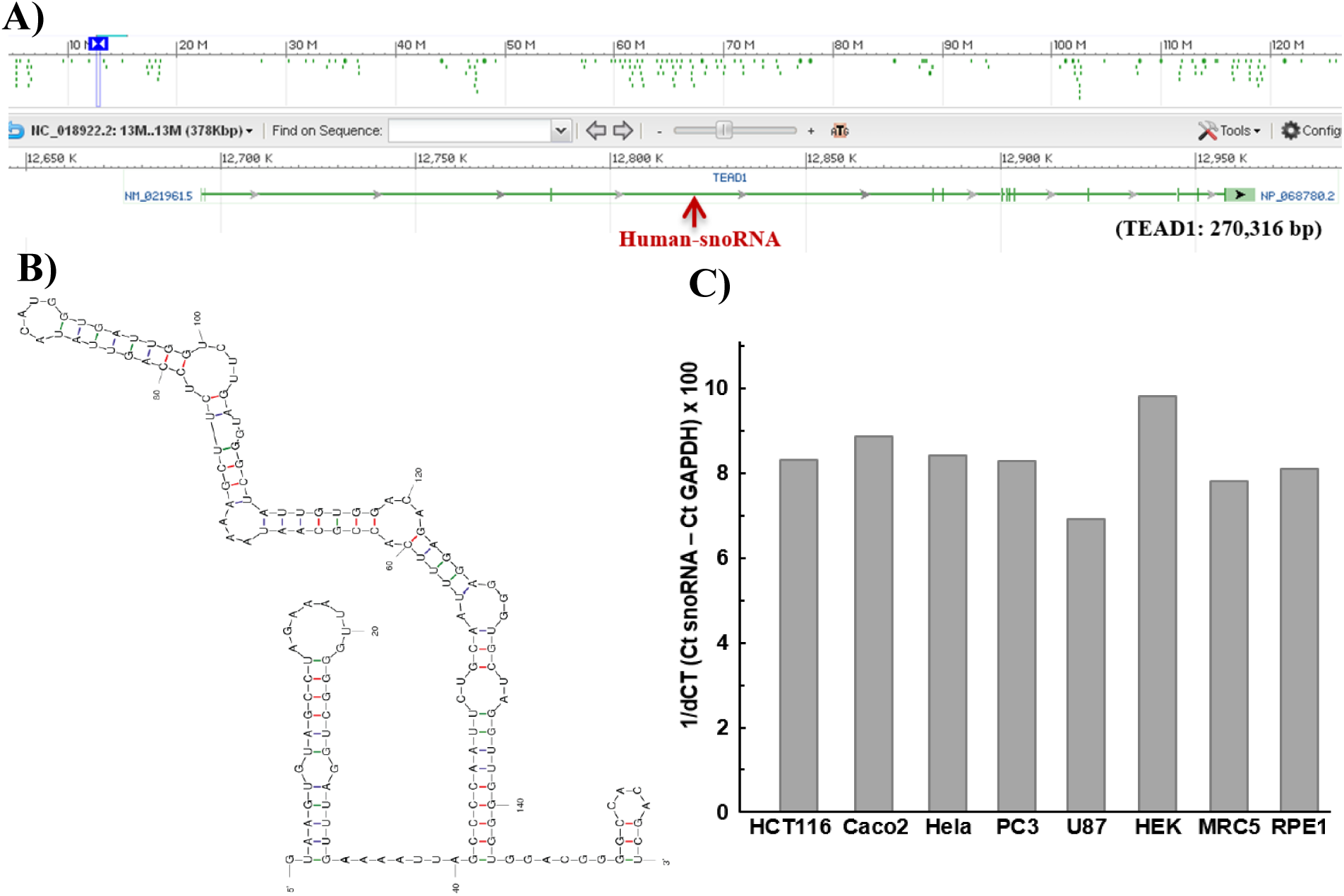
Genomic map, structure, and expression of the human snoRNA-*jouvence.* **A)** Genomic map of the snoRNA-*jouvence. h-jou* is localized on chromosome 11 (GRCh38.p13) in a large intron of the gene TEAD1. **B)** Secondary structure of the *h-jou*, as determined by the UNAFold software (http://unafold.rna.albany.edu/?q=DINAMelt/software). As predicted by its primary sequence, and like its *Drosophila* orthologue, *h-jou* is a H/ACA box forming a double hairpin. **C)** Expression level (1/dCt x 100) (dCT= Ct snoRNA - Ct GAPDH) in various cell lines determined by RT-qPCR (Taqman). Compared to the standard reference gene GAPDH, *h-jou* is weakly expressed in all different tested cell lines, with a very weak expression (to the limit of detection) in the U87 line (glioblastoma), and the strongest expression in the HEK293 cells (human embryonic kidney).

### The overexpression of *h-jou* stimulates the proliferation of the cells

In order to determine the effect of the *h-jou* on human cell lines, the human *jouvence* (159 bp) was stably transfected into HCT116, Caco2 and HEK293 cell lines respectively. The proliferation of these cells was compared to the corresponding vector (empty-plasmid) stably transfected cells. The proliferation was conducted for a period of nearly one week for the different cell lines. The cell counting was daily determined using the trypan blue exclusion with a VicellXR and/or by the quantification of the ATP with the CellTiter-Glo assay. For the 3 tested cells lines, the transfection of *jouvence* allowed cells to proliferate more rapidly. At 160 hours post-seeding, HCT116 *h-jou* transfected cells reached nearly 5 million cells *vs* only 2 million cells for the HCT116 empty plasmid cells: more than two folds (counted via the ViCellXR) (Figure 2A). The luminescence assay led to a similar difference (Figure 2B), although it is less pronounced. Similar result was observed for the HEK293-*jouvence* stably transfected cells compared to control cells (Figure 2C-2D) with an increase of about 50% of the cell number. Similarly, the Caco2 h-*jou* stably transfected cells were also more numerous than the corresponding empty plasmid cells (an increase of about 350 %) (Figure 2E). Therefore, we conclude that the overexpression of the *h-jou* importantly stimulates the proliferation of the stably transfected cells. In parallel, to support these results, the corresponding level of *h-jou* was assessed by RT-qPCR. HCT116 transfected cells were overexpressing *jou* for more than 300 fold compared to the vector (empty-plasmid) transfected cells (Figure 2F). Similarly, Caco-2 *h-jou* transfected cells were enriched with *jou* for nearly 200 fold (Figure 2G), while the HEK293 *h-jou* transfected cells had a fold-change of more than 1000 compared to the control cells (Figure 2H).

**Figure 2).**
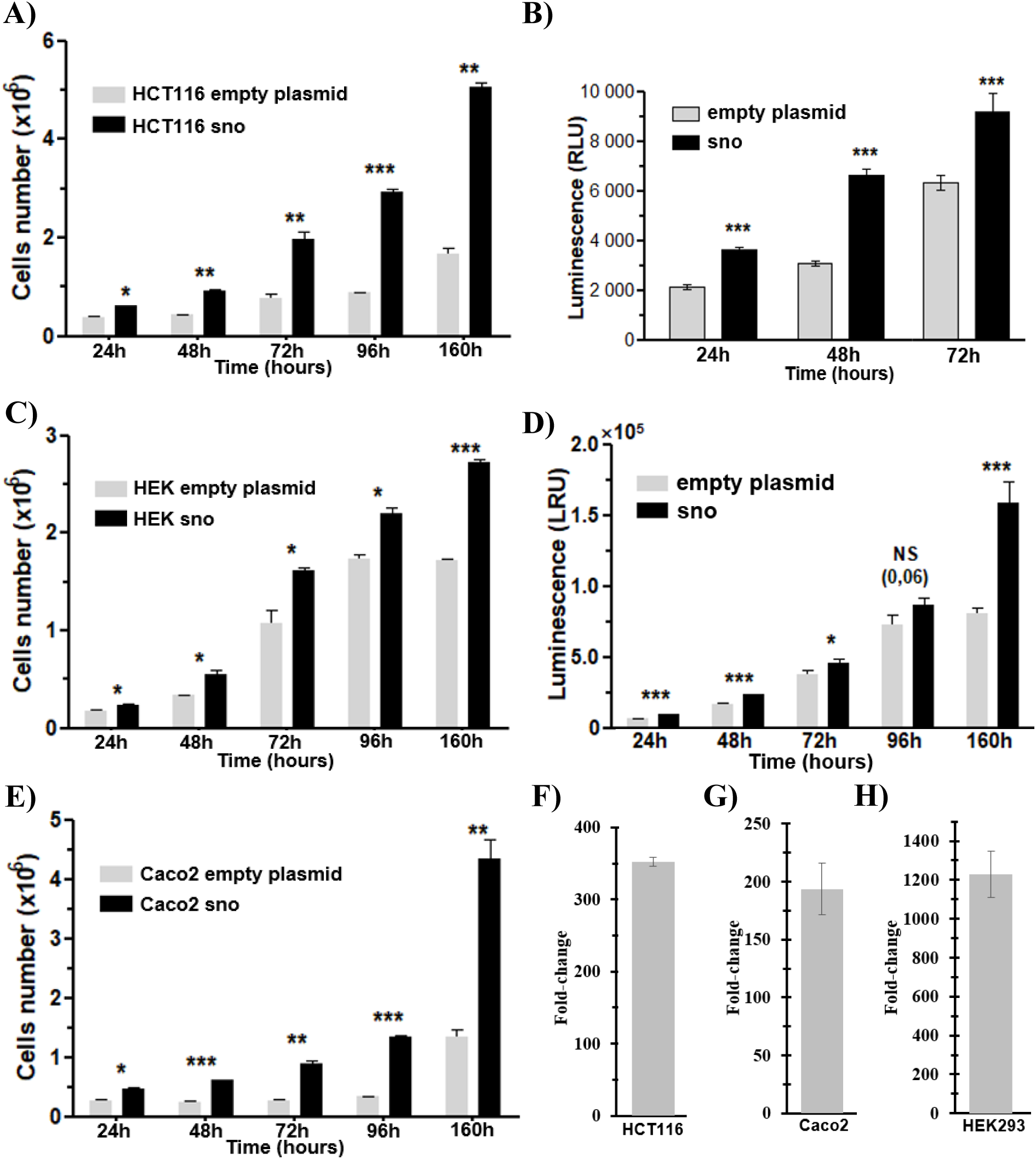
The overexpression of *h-jou* increases the proliferation of the cells. **A)** Cells number of HCT116 stably transfected cells with *jouvence* (in plasmid pcDNA-3.1) compared to the empty vector transfected cells determined by VicellXR. **B)** Same batch of cells than A, but spread and grow in 96 cell-plates and evaluated by luminescence (CellTiter-Glo), which quantify the amount of ATP, and consequently but indirectly, the number of cells. **C)** Cells number of HEK293 stably transfected cells with *jouvence* compared to the empty vector transfected cells determined by VicellXR. **D)** Same batch of cells than D, but spread and growth in 96 cell-plates and evaluated by luminescence (CellTiter-Glo). **E)** Cells number of Caco-2 stably transfected cells with *jouvence* compared to the empty vector transfected cells determined by VicellXR. The cells overexpressing *jouvence* are more numerous after 24 hours post-seeding, which have doubled their number after 72 hours, and again, more than tripled after 160h. **F-G-H)** Expression level (Fold change) of the transfected snoRNA-*jouvence* (overexpression) compared to non-transfected cells. **F)** In HCT116, *h-jou* is increased by about 350-fold. **G)** In Caco-2, the expression level of *h-jou* is increased by about 200-fold. **H)** In HEK293, the expression level is increased by about 1200-fold. Statistics: For the VicelXR (A-C-D) n=2, for the Luminescence CellTiter-Glo (B-D), n=6. Each figure is representative of three independent experiments. (p-values: * p<0,05; ** p<0,005; *** p<0,0005). Errors bars represent the mean +/- S.E.M. (p-value were calculated using the one-tail unpaired t-test using GraphPad Prism).

### Overexpression of *jou*vence by Lentivirus also increases the proliferation of the cell

As demonstrated above, the overexpression of *jouvence* leads to an increase of the proliferation of the cells. However, up to now, this effect has been observed only on stably transfected cell lines through the use of a plasmid and lipofectamine transfection. However, the constraints of this approach do not easily allow the overexpression of the *h-jou* in primary cells. In the aim to study the effect of the overexpression of *h-jou* directly on primary cells, we use the lentivirus approach. After building the lentivirus construct/vector and production of lentiviral particles (see Methods), we transduce the jou-lentivirus on various cells. First, we validate the lentivirus approach in HCT116 and CaCo-2 cells to compare the effect on proliferation to the previous observed data obtained in these cell lines. Using two different MOI (1 and 10), we observed, in HCT116 cells, an increase of cell proliferation determined by two independent readouts (Figure 3A-B), as previously described with the plasmid overexpression. In complement, we also measure by RT-qPCR the level of the *h-jou*. Interestingly we found roughly a 10-fold increase in the amount of *jouvence* (Figure 3C), with a slight increase according to the MOI concentration. In a step further, we also perform such transduction into the CaCo-2 cell line, and similarly, we observe an increase of their proliferation (Figure 3D-E). These similar results obtained on two independent cell lines validate the lentivirus approach, and consequently allow further exploring such approach on non-cancerous cell line and on primary cells. Then, transduction of *jou*-lentivirus on immortalized but non-cancerous RPE1 cell line (Human retinal pigmented epithelial cells) leads to a similar increase of cell proliferation (Figure 3F-G). Ultimately, the transduction the *jou*-lentivirus into primary cells (HUVEC) (Human Umbilical Vein Embryonic Cells) also leads to an increase of proliferation, as evaluated by the quantification of ATP using a luminescence assay (Figure 3H). In summary, the overexpression of *h-jou* using *jou*-lentivirus transduction system/vector yields to similar stimulation of the proliferation of non-cancerous primary cells.

**Figure 3).**
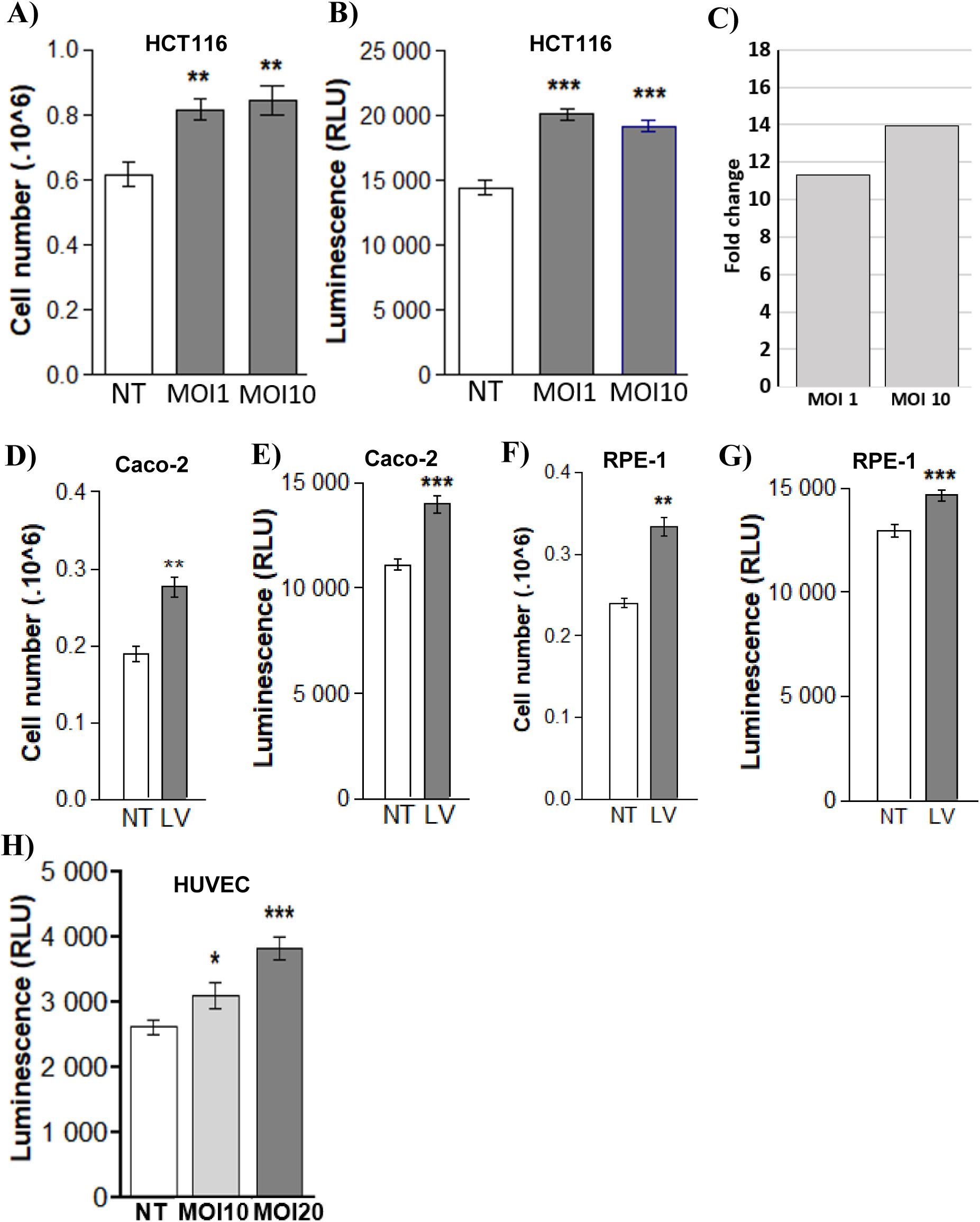
The overexpression of *jou* by lentivirus increases cells proliferation. **A)** Cells number of HCT116 cells transduced with *jou*-lentivirus, at two different MOI (1 and compared to non-transduced cells (NT) determined by VicellXR. **B)** Same batch of cells than A, but spread and grow in 96 cell-plates and evaluated by luminescence (CellTiter-Glo), which quantify the amount of ATP, and consequently but indirectly, the number of cells. Similarly, the amount of luminescence is increased after 96 hours for the two tested MOI. **C)** Expression level (Fold change) of the transduced *h-jou* (overexpression) compared to non-transduced cells. In HCT116, *h-jou* is increased by about 11-fold for the MOI-1, and 14-fold for MOI-10. **D)** Cells number of Caco-2 cells transduced with *jou*-lentivirus (LV), at MOI-10 compared to non-transduced cells (NT) determined by VicellXR. **E)** Same batch of cells than D, but spread and grow in 96 cell-plates and evaluated by luminescence (CellTiter-Glo). **F)** Cells number of RPE-1 cells transduced with *jou*-lentivirus (LV), at MOI-10 compared to non-transduced cells (NT) determined by VicellXR. **G)** Same batch of cells than F, but spread and grow in 96 cell-plates and evaluated by luminescence (CellTiter-Glo). **H)** Cells number of HUVEC transduced with *jou*-lentivirus (LV) at MOI-10 and MOI-20 compared to non-transduced cells (NT), spread and grow in 96 cell-plates and evaluated by luminescence (CellTiter-Glo). The amount of luminescence is increased after 96 hours. Statistics: For the VicelXR (A-D-F) n=3, for the Luminescence CellTiter-Glo (B-E-G-H), n=10. Each figure is representative of two independent experiments. (p-values: * p<0,05; ** p<0,005; *** p<0,0005). Errors bars represent the mean +/- S.E.M. (p-value were calculated using the one-tail unpaired t-test using GraphPad Prism).

### Decreasing *jouvence* level by siRNA reduces the proliferation of the cells

To assess if the decrease of the amount of *h-jou* could lead to a modified phenotype, the knockdown of *h-jou* was performed by transiently transfecting a *h-jou* specific siRNA to the HCT116 adenocarcinoma cell line. Briefly, a double siRNA transfection was performed: the first reverse transfection was made on HCT116 at the time of plating the cells into the culture plates, followed by a siRNA second transfection after 48 hours of culture. The number of cells was assessed after 72 or 96 hours depending of the tested cell lines. The negative siRNA mismatch control was also performed on the same conditions. The HCT116 transfected cells with jou-siRNA were proliferating in a lower rate compared to their two respective controls: the non-transfected cells, but treated with the lipofectamine/RNAiMax only (Co), or treated with the negative Si-RNA control (Si-Co), with the higher difference at 72 hours post-transfection (Figure 4A). The decrease of the amount of specific snoRNA by the siRNA was confirmed by RT-qPCR with a nearly 40% decrease of the endogenous *h-jou* at 72 hours post-transfection (Figure 4B). Therefore, we conclude that the partial inhibition (a decrease of 40%) of the *h-jou* was sufficient to decrease the proliferation of the cells. To assess if this cellular effect of the decrease of the *h-jou* is not cell type specific, we perform similar experiments on other well-established cell lines. Similarly, striking decreases of the cell number were observed for the MCF7 cells (a breast cancer cell line) (Figures 4C-D), the U87-MG cells (a glioblastoma cell line) (Figures 4E-F), and the A549 lung cancer cell line when the *h-jou* is knocked-down (Figures 4G-H). Finally, since the four cell types investigated above are all immortalized cancerous cells, we wondered if similar effects can be observed in primary non-cancerous cells. The *h-jou* knock-down on HUVEC induces a similar decrease of cell proliferation determined both by cell counting ViCellXR (Figure 4I), and by ATP measurement (CellTiter-Glo) (Figure 4J). These similar results obtained in several different cancerous cell lines as well as on primary cells indicate that a normal and physiological level of *h-jou* is required for the proper cell proliferation. Indeed, increasing the level of *h-jou* stimulates the proliferation, while inversely decreasing its level by siRNA inhibits the proliferation of the cells. It also indicates that *h-jou* is functional in all these cell lines derived from different organs and tissues, although *h-jou* is only expressed at a low level (see Figure 1C).

**Figure 4).**
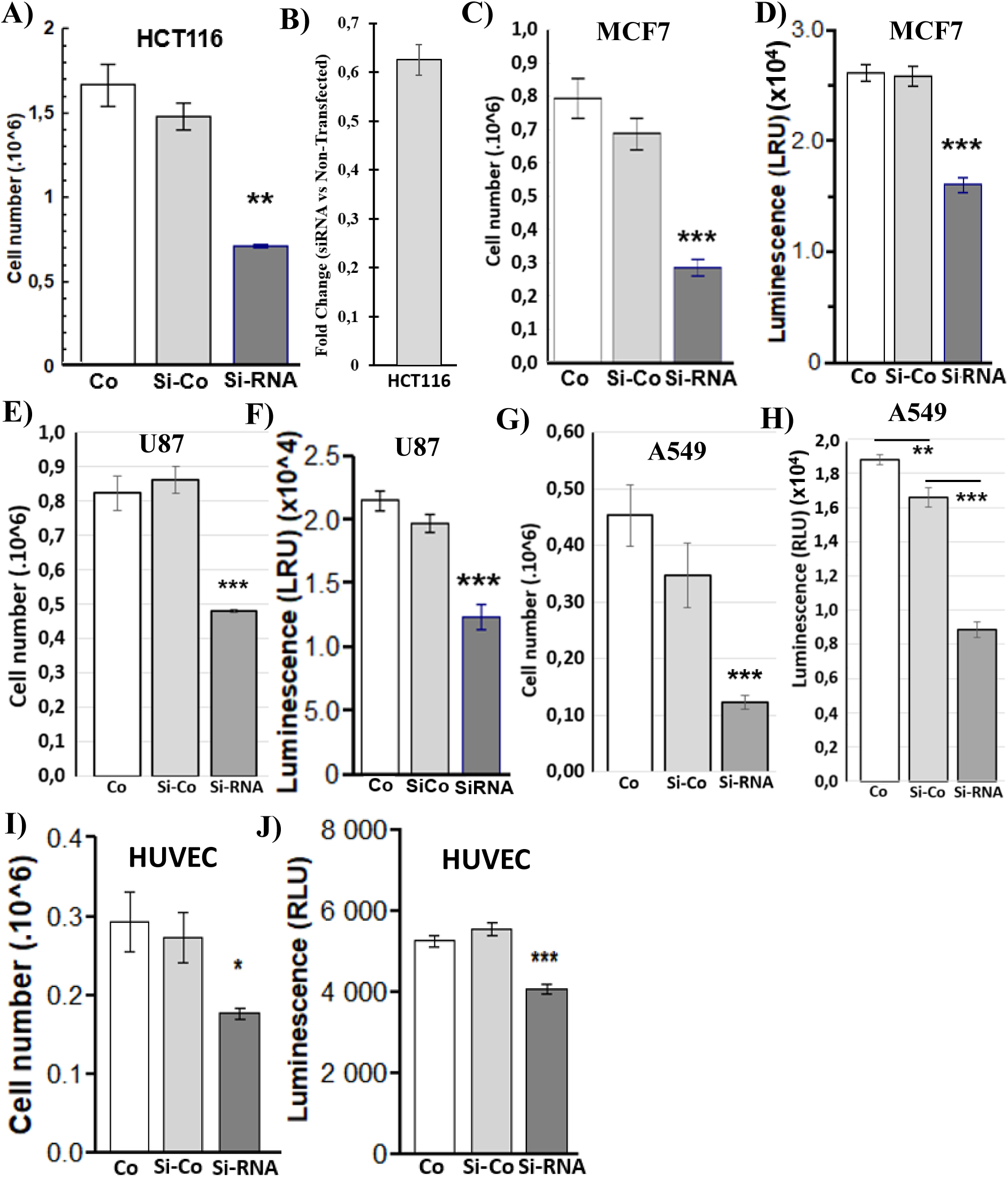
The knockdown of *h-jou* by si-RNA decreases the proliferation. **A)** Cells number of HCT116 transiently transfected cells with LNA-siRNA directed against *jouvence* compared to the control non-transfected cells (Co = treated with the same amount of RNAiMAX), and to the control (Si-Co = transfected cells with a siRNA-control without target), determined by VicellXR 72 hours post-seeding. **B)** The expression level (Fold change) of the siRNA transfected HCT116 cells compared to the non-transfected cells, determined 72 hours post-seeding (same batch of cells than A). **C)** Cells number of MCF7 transiently transfected cells compared to their two respective controls, as in A, determined by VicellXR. **D)** Same batch of cells than C, but spread and growth in 96 cell-plates and evaluated by luminescence (CellTiter-Glo) after 96 hours. **E)** Cells number of U87 transiently transfected cells with LNA-siRNA compared to their respective controls, determined by VicellXR. **F)** Same batch of cells than E, but spread and growth in 96 cell-plates and evaluated by luminescence after 96 hours. **G)** Cells number of A549 transiently transfected cells with LNA-siRNA compared to their control, determined by VicellXR. **H)** Same batch of cells than G, but spread and growth in 96 cell-plates and evaluated by luminescence after 96 hours. **I)** Cells number of HUVEC (primary cells) transiently transfected cells with LNA-siRNA compared to their respective control, determined by VicellXR. **J)** Same batch of cells than I, but spread and growth in 96 cell-plates and evaluated by luminescence. Statistics: For the VicelXR (A-C-E-G-I) n=3, for the Luminescence CellTiter-Glo (B-D: n=27), (H: n=8), (J: n=10). Each figure is representative of three independent experiments. (p-values: * p<0,05; ** p<0,005; *** p<0,0005). Errors bars represent the mean +/- S.E.M. (p-value were calculated using the one-tail unpaired t-test using GraphPad Prism).

### HCT116 overexpressing *h-jou* presents a genomic signature of dedifferentiation

In order to initiate the elucidation of the genetic and molecular mechanisms of the snoRNA-*jouvence* that could be responsible for these phenotypes, we have characterized the cells on a whole transcriptomic level. A RNA-seq analysis was performed on HCT116 cells, comparing the *h-jou* stably transfected cells to their empty plasmid control. The comparison reveals a set of 5918 Differentially Expressed Genes (DEG), with 2974 up-regulated genes and 2944 down-regulated genes (Figure 5A). Based on the fixed p-value, Table 1A shows the list of 19 most up-regulated genes (for the complete list of the up-regulated genes, see the Suppl. Table S1), while the Table 1B shows the list of 19 most down-regulated genes (see Suppl. Table S2 for the complete list of down-regulated genes). The cluster analysis (Heat-map) of differentially expressed genes briefly resumes the up- and down-regulated genes (Figure 5B). To get more precise information about these multiple deregulated genes, the statistical enrichment of DEG in KEGG pathway [14,15] shows an important enrichment in metabolic pathways, with more than 400 DEG with a relatively low Rich Factor but with a high q-value (adjusted p-value) (Figure 5C). Other pathways were also enriched (with high q values), such as RAS signaling pathway, PI3K-AKT pathway, regulation of actin cytoskeleton pathway, pathways in cancer, and the ribosome pathway having the higher degree of enrichment, with a Rich Factor of more than 0,7. Again here, based on the p-value, Table 1C shows the list of the 18 most affected KEGG pathways (see Suppl. Table S3 for the complete list of affected KEGG pathways).

**Table 1).**
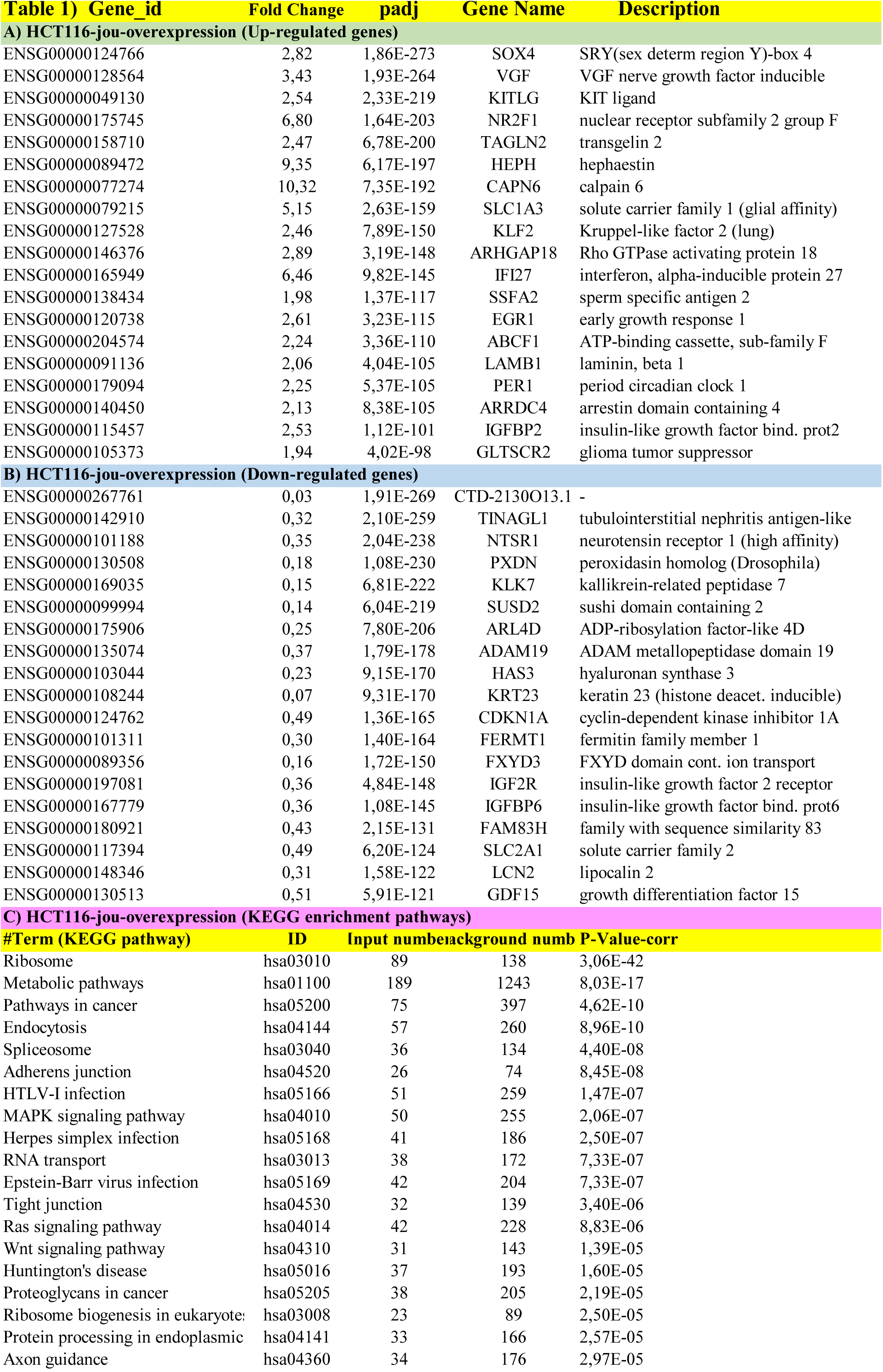
Short list of the main genes deregulated in overexpressing *jouvence* HCT116 cells revealed by RNA-Seq. **A)** Up-regulated genes. **B)** Down-regulated genes. **C)** Short list of the main deregulated pathways revealed by the KEGG analysis. For the complete list, see Suppl. Tables S1-S2-S3 respectively.

**Figure 5).**
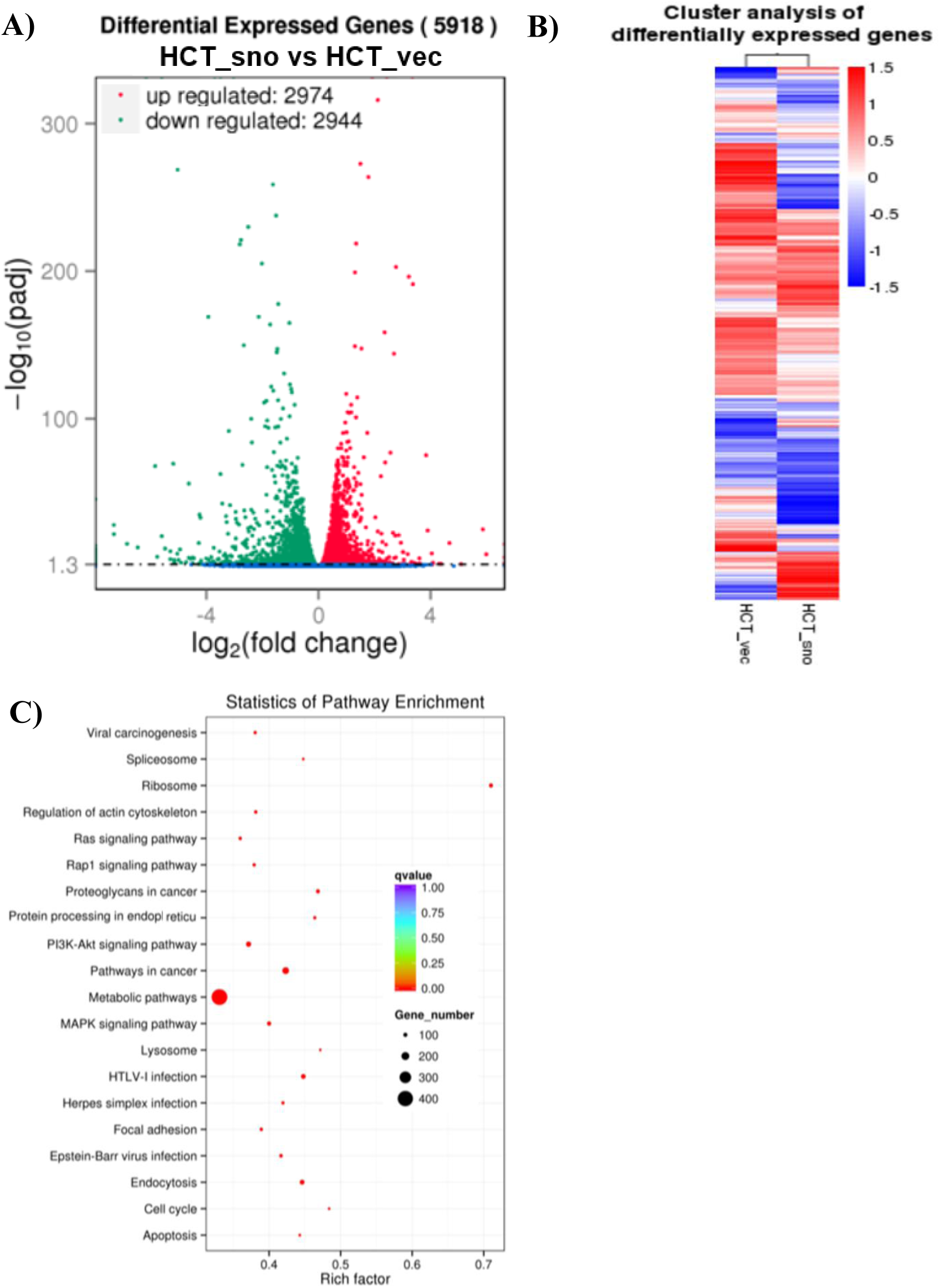
Hundreds of genes are deregulated in HCT116 cells overexpressing *jouvence*. **A)** Transcriptomic analysis (RNA-Seq) performed on total-RNA from the overexpressing *jouvence* HCT116 cells compared to the empty vector cells, reveals that 5918 genes are deregulated, in which 2974 are upregulated, while 2944 are downregulated (see Suppl. Table-S1 and Suppl. Table-S2 for the complete list of genes). **B)** Clusters analysis (Heat map) of differentially expressed genes. **C)** Statistic of enrichment pathway of the deregulated genes according to the KEGG analysis (see Suppl. Table S3) for the full list of KEGG analysis. In brief, the metabolic pathways are the main deregulated pathways in term of number of genes, while the ribosome is the main deregulated pathway in term of strength (Rich factor).

More specifically, among the 2974 up-regulated genes, an important number of them harboring the best adjusted p-values are found to be correlated with a dedifferentiation of the cells [16]. Notably, these genes are up-regulated, as in an EMT (Epithelial-Mesenchymal Transition) context, which is controlled by four major interconnected regulatory networks [17-19]. For examples, among them, the VGF nerve growth factor inducible (VGF) known to play a role in the cell plasticity and to induce the transcription factor TWIST1, which facilitate EMT in cancer cells [20]. The Kit Ligand (KITLG), a ligand for the receptor-type protein-tyrosine kinase plays an essential role in cell survival, proliferation, hematopoiesis and stem cell maintenance [21]. The nuclear receptor subfamily 2, group F, member 1 (NR2F1), is a nuclear hormone receptor and a transcriptional regulator. It is associated with stem cells, and acquisition of stem-like properties and quiescence [22]. Transgelin 2 (TAGLN2), an actin-binding protein, is significantly induced in hypoxic cancer cells, while Snail1 is simultaneously increased, and thus, inducing the EMT by downregulating E-cadherin [21,23]. The solute carrier family 1 (glial high affinity glutamate transporter), member 3 (SLC1A3), a glutamate transporter mediates the inter-niche stem cell activation [24]. The early growth response 1 (EGR1) factor has a role in controlling cell plasticity, which has been involved in TGFβ 1-induced EMT [25]. The Kruppel-like factor 2 (KLF2) family are considered as key transcription factors implicated in self-renewal of embryonic stem cells [26]. All of these key genes, among others, are considered as landmark of EMT.

Inversely, 2944 genes are under-expressed. Interestingly, also here, an important number of these down-regulated genes have been shown to be deregulated during the EMT, which again suggests that the overexpression of *h-jou* induces EMT. For examples, the Tubulointerstitial nephritis antigen-like 1 (TINAGL1) decreases the secretion of metastasis-suppressive proteins including insulin-like growth factor binding protein 4 (IGFBP4) [27]. The hepatocyte nuclear factor 4α (HNF4A) is under-expressed (fold-change 0,02), while the EMT in hepatocytes correlates with the down-regulation of the hepatic differentiation key factors HNFs [28]. The Keratin 13 (KRT13) is epigenetically suppressed during Transforming Growth Factor-β 1-induced epithelial-mesenchymal transition [29]. The catenin (cadherin-associated protein), alpha-like 1 (CTNNAL1), an epithelial marker is also under-expressed [30]. The Myosin binding protein H (MYBPH) which inhibits cell motility and metastasis, and then linked to an EMT phenotype, is also under-expressed [31]. In addition Claudin 9 (CLDN9), an epithelial marker [32] is under-regulated in the *h-jou* overexpressing cells, while it is found to be overexpressed in the siRNA transfected HCT116 cells (see below). Annexin 8 (ANXA8) is down-regulated (with a fold-change of 0,1). This last has been shown to be transcriptionally down-regulated by epidermal growth factor (EGF), which correlates with the morphologic changes of the epithelial-to-mesenchymal transition (EMT) [33], along with tumor dedifferentiation. To finish the selected list of markers, the epithelial marker Protocadherin 1 (PCDH1) is also down-regulated. PCDH1 binds to SMAD3 and suppresses TGFβ1-induced gene transcription [34].

In addition, the overexpressing *h-jou* transfected cells are also enriched in CSC (Cancer Stem Cells) potential markers, such as : SOX4, SOX8, CD44, MSI-2, EpCAM to name but of few (see Figure 7). Sox family expression has been correlated with mesenchymal traits and loss of epithelial features. SOX4 plays a role in TGFβ-induced EMT and confers stem cell characteristics [35]. SOX8 regulates CSC properties and EMT via Wnt β-catenin pathway [36]. The cell surface antigen CD44, Musashi RNA-binding protein 2 (MSI-2) and the Epithelial Cell Adhesion molecule (EpCAM) are also potential CSC markers [37], and are found to be overexpressed in *h-jou* transfected HCT116 cells. In addition, an important number of KEGG pathways are significantly deregulated (Figure 5C and Suppl. Table S3), as the Insulin Secretion Pathway, and the Insulin signaling pathway. Many of the lipid metabolic pathways are also deregulated, such as Glycerophospholipid pathway, Glycerolipid metabolism, sphingolipid metabolism pathway. The Longevity regulating pathways are also affected, with genes like IRS2, AKT3, IGF1R, NFkB1, FOXO1, and RPTOR. These pathways are linked (directly or indirectly) to the EMT, the plasticity and the longevity of the cells. Finally, to support the RNA-Seq analysis, we also validated, by RT-qPCR some of the DEG (Suppl. Figure 1A).

**Figure 6).**
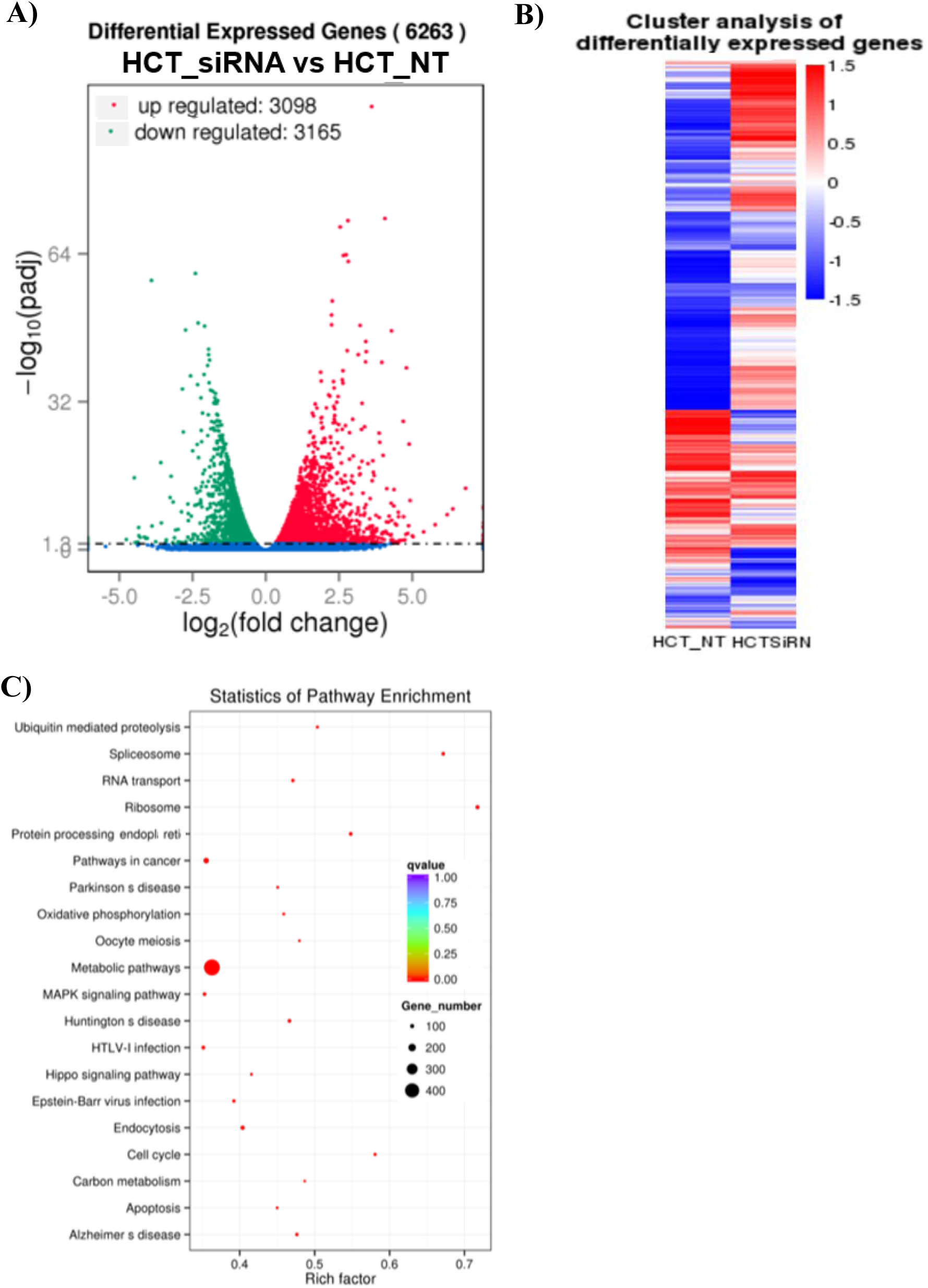
Hundreds of genes are deregulated in *jouvence* depleted HCT116 cells. **A)** Transcriptomic analysis (RNA-Seq) performed on total-RNA from the knockdown of *jouvence* by siRNA on HCT116 cells compared to the control non-transfected cells, reveals that 6263 genes are deregulated, in which 3098 are upregulated, while 3165 are downregulated (see Suppl. Table-S4 and Suppl. Table-S5 for the complete list of genes). **B)** Clusters analysis (Heat map) of differentially expressed genes. **C)** Statistic of pathway enrichment of the deregulated genes according to the KEGG analysis (see Suppl. Tables S6 for the full list of the KEGG analysis). In brief, the metabolic pathways are the main deregulated pathways in term of number of genes, while the ribosome and spliceosome are the two main deregulated pathways in term of strength (Rich factor).

**Figure 7).**
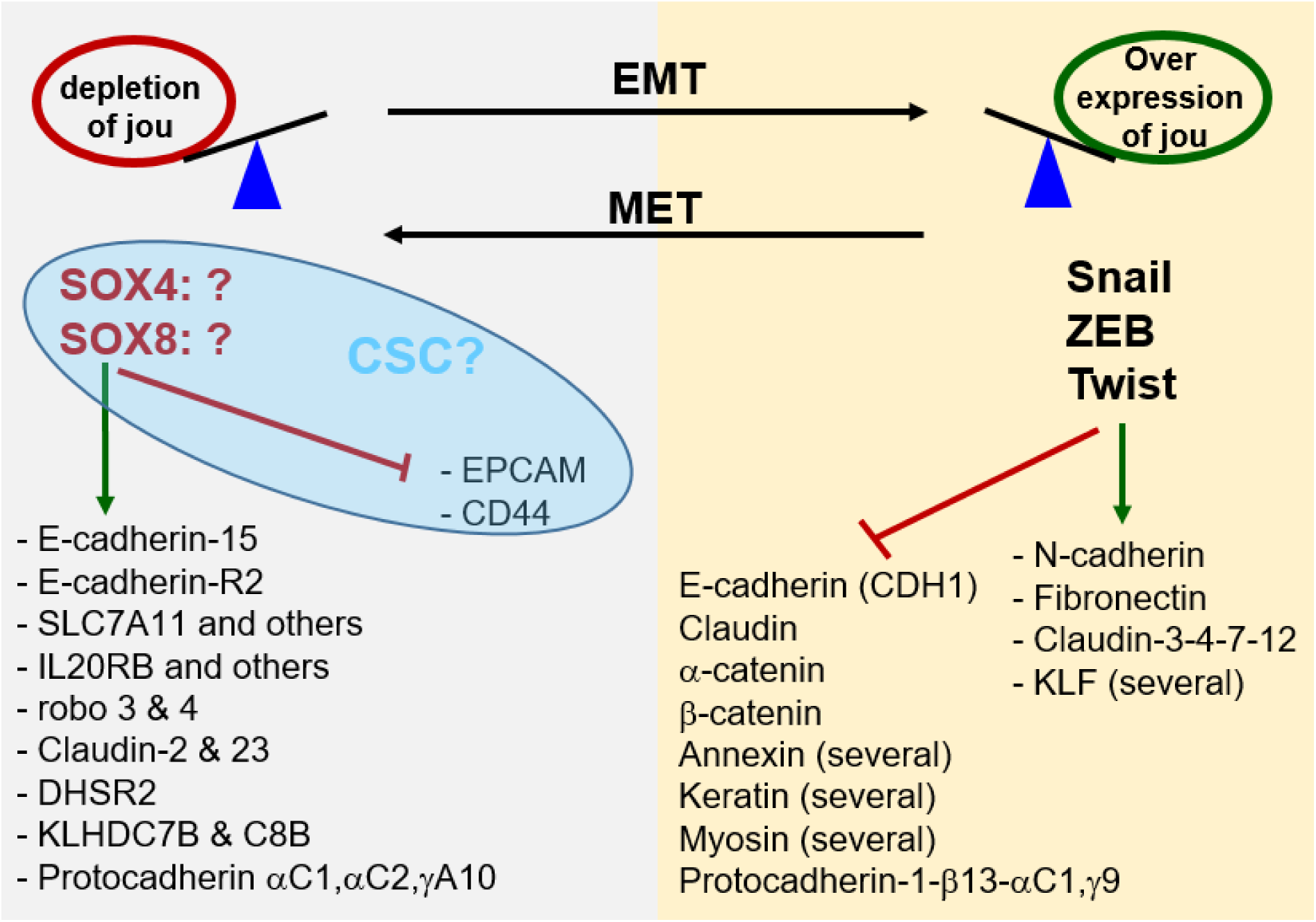
Summary of main deregulated genes suggesting a dedifferentiation. Several of the main key Transcription Factors genes (Twist, Snail, ZEB) involved in EMT are upregulated in *jou*-overexpression. Consequently, several of their known targets as N-cadherin, Fibronectin, few Claudin and several KLF are upregulated. In parallel several genes are downregulated, as E-cadherin (CDH1), Claudin, α-catenin, β-catenin, several Annexin, several Keratin and several Myosin, as well as Protocadherin1-β13, αC1, γ9. Inversely, in the *jou-* depleted cells by siRNA, we rather observe the opposite phenotype, suggesting a MET (Mesenchymal-Epithelial-Transition). More particularly, we observe a decrease of two Transcription Factors as SOX4 and SOX8, although these two last have not yet been clearly demonstrated to induce MET. Several other genes (putatively their targets) are upregulated, while two key genes, as EPCAM and CD44 are downregulated. This last group of four genes (blue ellipse) could also suggests a CSC phenotype. Nevertheless, although we do not observe all the characteristic cells markers of the EMT, several of the deregulated genes in *jou*-overexpression strongly suggest an EMT, or at least a partial or hybrid EMT, as suggested by Pastushenko and Blanpain (2019) [18], while inversely, the knock-down of *jouvence* rather seems to direct the cells toward a MT.

### Knockdown of *h-jou* by siRNA decreases the ribosome biogenesis and spliceosome

Similarly, a RNA-seq analysis was performed on HCT116 cells with a depletion of *jouvence*. The comparison of the HCT116 *h-jou* knocked-down by siRNA transfected cells compared to the HCT116 non-transfected cells (treated with lipofectamine/RNAiMax only) reveals a set of 6263 Differentially Expressed Genes (DEG), with 3098 up-regulated genes and 3165 down-regulated genes (Figure 6A). The Table 2A shows the list of 19 most up-regulated genes (for the complete list of the up-regulated genes, see the Suppl. Table S4). The cluster analysis of differentially expressed genes briefly resumes the up- and down-regulated genes (Figure 6B). KEGG analysis pathways enrichment presents an enrichment in the Metabolic pathways, with the highest number of DEG (∼ 400) (Figure 6C). However, the Spliceosome and Ribosome KEGG pathways had the highest Rich Factor (around 0,7). Pathways like RNA transport, Ubiquitin Mediated proteolysis, Protein processing in the endoplasmic reticulum, Pathways in cancer, MAPK signaling pathway, Hippo pathway, to name but a few, were also enriched with low q values (around 0) and a Rich Factors varying from 0,4 to nearly 0,6.

**Table 2).**
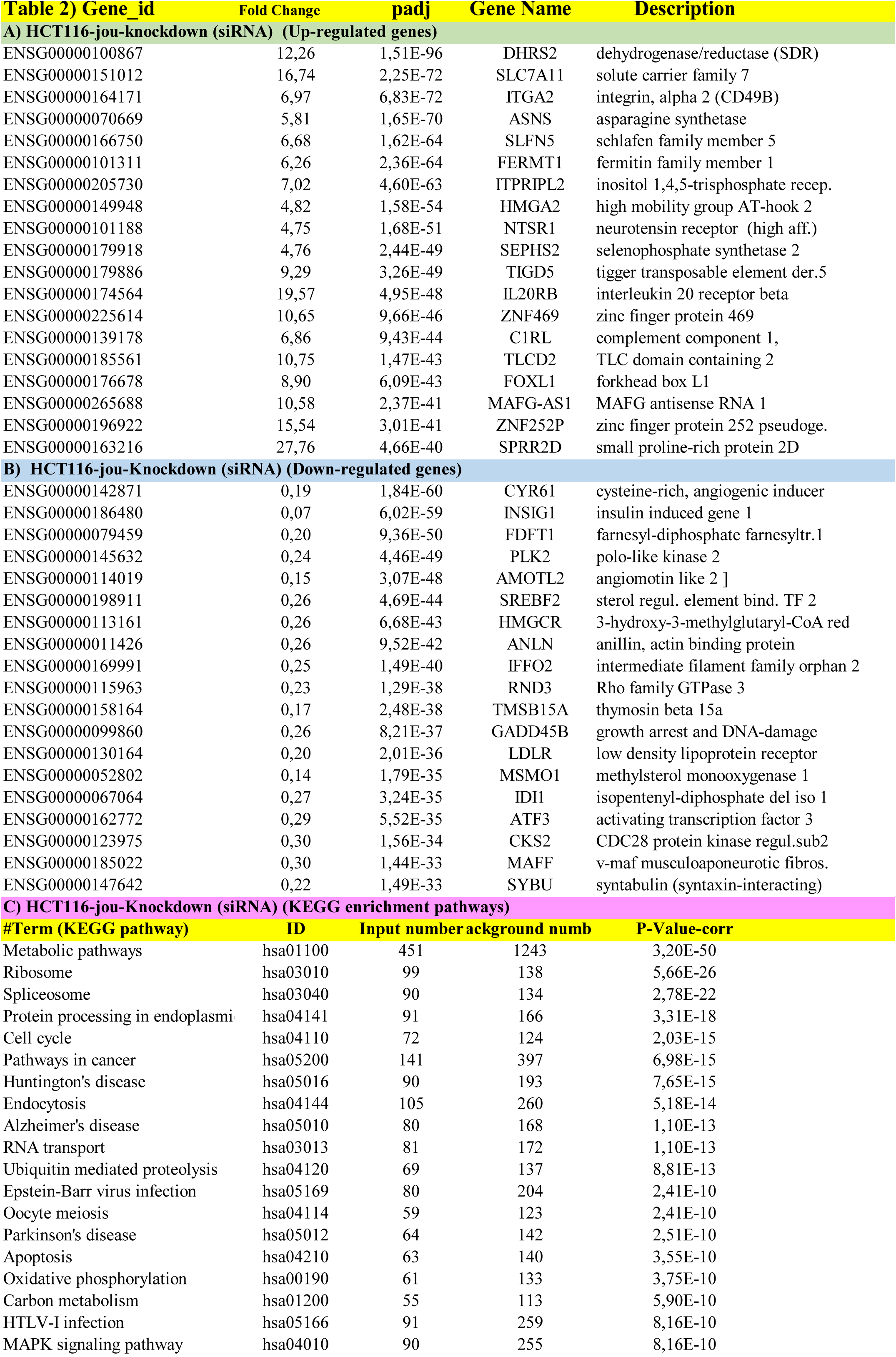
Short list of the main genes deregulated in knockdown *jouvence* HCT116 cells revealed by RNA-Seq. **A)** Up-regulated genes. **B)** Down-regulated genes. **C)** The main deregulated pathways revealed by the KEGG analysis. For the complete list, see Suppl. Tables S4-S5-S6 respectively.

More specifically, the Solute Carrier family 7 (anionic amino acid transporter light chain, xc-system), member 11 (SLC7A11) and interleukin 20 receptor-β (IL20RB) are upregulated in the siRNA condition. Kelch domain containing 7B (KLHDC7B), which has been associated with gene modulation activity in the Interferon signaling pathway [38], is also upregulated after *h-jou* knock-down (fold-change of 83,3). Roundabout, axon guidance receptor, homolog 4 Drosophila (ROBO4) which increases cell adhesion [39] and correlates with an epithelial phenotype, is upregulated (with a fold-change of 73,09). Moreover, Cadherin 15, type 1, M-cadherin (CDH15) and Claudin 2 (CLDN2), which are epithelial markers, are overexpressed when *h-jou* is depleted. This suggests that the inhibition of the snoRNA could lead to more pronounced epithelial phenotype (MET) while inversely, its overexpression favors the EMT.

On the other hand, the knockdown of *h-jou* leads to the down-regulation of 3165, with a majority of down-regulated genes belonging the ribosome or spliceosome pathways. Table 2B shows the list of 19 most down-regulated genes (for the complete list of the down-regulated genes, see the Suppl. Table S5). This refers to the implication of the H/ACA box snoRNAs in the ribosome biogenesis, the modification and processing of ribosomal RNA precursors [1-4]. More specifically, splicing factors are down-regulated such as serine/arginine-rich splicing factor 4 (SRSF4), serine/arginine-rich splicing factor 11 (SRSF11), splicing factor 3b, subunit 1, 155kDa (SF3B1) and splicing factor 3b, subunit 2, 145kDa (SF3B2) (Suppl. Table S5). Some ribonucleoproteins are also down-regulated: heterogeneous nuclear ribonucleoprotein H1 (HNRNPH1), small nuclear ribonucleoprotein 70kDa (U1) (SNRNP70), small nuclear ribonucleoprotein polypeptide E (SNRPE) (Table 2B and Suppl. Table S5). In other words, the main deregulated KEGG pathways (with the highest Rich Factors) are the Ribosome and the Spliceosome pathways (Table 2C). More strikingly, the depletion of *h-jou* leads to the down-deregulation of the majority of the genes of these pathways. For the Ribosome pathway, 90 genes were down-regulated over the 99 genes. For the Spliceosome pathway, while the total number of genes is 90, among them, 80 were down-regulated. These effects perfectly correlate with the main known function of the snoRNAs in guiding chemical modifications of other RNAs, mainly rRNAs, transfer RNAs and small nuclear RNAs, through their function within their related RNPs. Finally, we have also validated, by RT-qPCR, some of these DEG: CLN2, HMGCR, PRPF3, SRSF4, DHRS2, IL20RB, SLC7A11 (Suppl. Figure 1B).

### Several genes are deregulated in opposite direction when comparing overexpression and depletion of *jou*

The RNA-Seq analysis has revealed that several genes are differentially expressed (DEG) either up or down, and so, both in overexpression or in knockdown of *h-jou*, related to a significant number of enriched KEGG pathways. Then, we wonder if the genes that are upregulated in *h-jou* overexpression condition are inversely downregulated in the inverse *jouvence*-knocked-down condition, and vice-versa. Thus, when the upregulated genes in the *jouvence*-overexpression condition were compared to the down-regulated genes in the *jouvence* depletion (siRNA condition), we found 1102 DEG in common (see Suppl. Table S7 for the list of genes). Conversely, the comparison of the down-regulated genes in *jouvence* overexpression condition versus the overexpressed genes in *jouvence* depletion (siRNA condition) led to a number of 868 DEG in common (see Suppl. Table S8 for the list of genes). For examples, among the 3098 overexpressed genes, the genes DHRS2, SLC7A11, IL20RB, KLHDC7B, ROBO4, CDH15, CDLN2, that are up-regulated when the *h-jou* is inhibited were found to be down-regulated in the inverse condition of *h-jou* stably transfected cells (overexpression). Briefly, Dehydrogenase/reductase (SDR family) member 2 (DHRS2) which inhibits cell growth and motility [40], was overexpressed in the siRNA cells, which correlates with the observed phenotype after siRNA transfection, where proliferation rate decreased significantly at 72 hours post-transfection-1. Interestingly, these results indicate that *h-jou* leads to the up- or down-regulation of a similar set of genes depending of its over- or under-expression. Consequently, it suggests that these multiple genes are directly sensitive to the level of *h-jou*, and therefore are likely not only a consequence of the deregulation of other genes.

## Discussion

We have characterized a new snoRNA named *jouvence* in human, which was not yet annotated in the genome. *jouvence* is localized on the chromosome 11 in a large intron of the gene TEAD1. As previously reported [13], in addition to some tissues, *h-jou* is expressed, although weakly, in all tested various well-established cell lines. To gain insights about the cellular role of *jouvence*, we first overexpress *h-jou* into different cancerous and non-cancerous cell lines. Interestingly, the overexpression of *jouvence* leads to an important increase of the proliferation of the cells, yielding to about twice more cells within about one week. More importantly, this effect is observed with two independent approaches to overexpress *h-jou*, first through a stably transfected-plasmid and second through a transduction with a lentivirus vector. This second result obtained with an integrative lentivirus indicates that a snoRNA can be successfully carried by lentivirus. Inversely, the decrease (knockdown) of *h-jou* by transiently transfected specific siRNA, leads to an opposite phenotype characterized by a rapid decrease of cell proliferation.

To investigate further the role of the *h-jou* and more particularly its molecular mechanism, we perform a transcriptomic analysis (RNA-Seq). The overexpression of the *h-jou* leads to the up- and down-regulation of many genes. Among them, several deregulated genes, as Twist, SNAIL/SMUG/SMUC, ZEB, Fibonectin, suggest a dedifferentiation signature of the cell (Figure 7). This cellular effect could be comparable, at least in part, to a rejuvenation of the cells [41]. It could also remind, in some ways and in a larger context, the organismal phenotype observed in *Drosophila*, an extension of lifespan [13]. On the other hand, the RNA-Seq performed on the decrease of *h-jou* through the knockdown by siRNA, also shown that several genes are deregulated, both up and down, while the bioinformatics analysis has revealed that the main affected KEGG pathways are, in addition to the metabolic pathway, the ribosomes biogenesis and the spliceosome. These results are in agreement with the canonical role of a H/ACA box snoRNA, known to be involved in the regulation of the ribosome biogenesis and the modification of the ribosomal RNA [1-4]. More precisely, according to the KEGG pathways, 99/138 genes of the ribosome pathway are affected, and among them, 91/99 are downregulated. For the spliceosome pathway, 90/134 genes are affected, and among them, 80/90 are downregulated. In overview, these results indicate a clear and an almost complete breakdown of the ribosome and the spliceosome pathways, which consequently might explain the decrease of cell proliferation. It has been reported that repressing or perturbing the ribosome biogenesis decreases the growth of the cells [42], and/or even could drive cells into senescence [43].

The Epithelial to Mesenchymal Transition (EMT) is a process by which epithelial cells lose cell-cell adhesion, and gain migratory and invasive properties to become mesenchymal stem cells [18,19,21]. The cancer cell lines with mesenchymal phenotype are characterized by under-expression of different genes such as claudins, cadherins, occludins, among others. In addition, genes such as Fibronectin, Jagged 1 are overexpressed, along with the three main EMT transcriptional factors : SNAIL/SLUG/SMUC, Twist and ZEB [17]. These factors can act together to induce EMT. Indeed, human SNAIL1 (SNAI1) protein encoded by SNAI1/SNA gene represses the transcription of E-cadherin/CDH1 gene (Figure 7). Human SNAIL2 (SNAI2) protein encoded by SNAI2/SLUG gene induces the first phase of epithelial-mesenchymal transition (EMT), including desmosome dissociation, cell spreading, and initiation of cell separation [17]. Twist1 activates other EMT-inducing transcription factors to suppress E-cadherin and promote EMT and tumor metastasis. Twist1 needs to induce Snail2 to suppress the epithelial branch of the EMT program, while Twist1 and Snail2 act together to promote EMT [17]. ZEB is regulated by multiple signaling pathways, such as WNT, NF-κB, TGF-β, COX2, HIF, among others [17,44].

Remarkably, the overexpression of the new H/ACA *h-jou* seems to be sufficient by itself to re-orientate the cells toward the dedifferentiation, although the HCT116 cells (an adenocarcinoma line from the gut) is originally cancerous cells. Nevertheless, the overexpression of *h-jou* seems to superimpose a signature of dedifferentiation. Indeed, several key genes involved in the EMT (Epithelial-Mesenchymal Transition), as TWIST, SNAIL/SMUG/SMUC, ZEB, Jagged and fibronectin, are upregulated, while some of their target genes such as claudin, E-cadherin, annexin, and α-catenin are downregulated (Figure 7) [19,21,45]. Nonetheless and alternatively, at this stage of this study, we could not exclude that this EMT process could rather be interpreted as a step toward the acquisition of a cancer stem cell (CSC) phenotype. Indeed, some evidence suggests that snoRNAs may also play a role in cancer stem cells (CSC) [12]. For example, ALDH1 has been demonstrated to be a CSC marker [46]. Here, several deregulated genes could also suggest a CSC mechanism, as for instance, few ALDH1-subunits, as ALDH1A3, ALDH1B1, and ALDH1L2. Other genes as SOX4, SOX8, CD44, MSI-2 and EPCAM, which are also deregulated in *h-jou* depletion have also been considered as potential CSC markers [47] (Figure 7). Further experiments will be required to discern between a CSC trend rather than an EMT dedifferentiation trend.

Based on its primary sequence, *jouvence* is a snoRNA of type H/ACA box, generally known to perform pseudouridylation [1]. Pseudouridylation is a post-transcriptional isomerization reaction that converts uridine to pseudouridine (Ψ) [48]. This last is present in several different types of RNAs, including coding and noncoding RNAs [5,49]. Ψ is particularly concentrated in rRNA, in which it plays an important role in protein translation, as well as in spliceosome snRNAs, in which it is involved in pre-mRNA splicing. Here, disrupting the level of *h-jou*, either by overexpression or by decreasing its expression, leads to the deregulation of a quite huge number of genes (about 6000: about half up and half down), as revealed by the RNA-seq analysis. Interestingly, among them, about 1000 genes (868 and 1102 respectively) are deregulated in the opposite direction when we compare *h-jou* overexpression with its depletion by siRNA (or vice-versa), suggesting that the expression level of these genes follows directly the level of *jou*, and consequently might be likely directly regulated by the snoRNA-*jouvence*. In counterpart, it suggests that the remaining deregulated genes (∼ 5000) are rather likely a consequence of the deregulation of these “primary” 1000 genes, or in another term, a secondary effect. Further experiments will be required to decipher the precise molecular mechanisms of *jouvence* which leads to the deregulation of these multiple genes, either due to transcription or translation modifications, chromatin remodelling, RNA stability, or else.

In addition to the well-known canonical function of the snoRNA in various RNA modifications, more recently, some snoRNAs have been involved in stress response and metabolic homeostasis. For examples, the snoRNA U32a, U33 and U35a are mediators of oxidative stress [50], while the same snoRNAs seem also to regulate the systemic glucose metabolism [51]. In the same way, Brandis et al. [52] has shown that the box C/D snoRNA U60 regulates intracellular cholesterol trafficking between the plasma membrane and the endoplasmic reticulum. Similarly, the snoRNA U17 regulates the cellular cholesterol trafficking through the hypoxia-upregulated mitochondrial regulator (HUMMR) by acting on its target mRNA [53]. Here, the modification of the level of the *h-jou*, either its overexpression or its depletion leads to a strong deregulation of metabolic pathways as revealed by the KEGG analysis (about 400 genes). Interestingly, among them, the 3-hydroxy-3-methylglutaryl-CoA reductase (HMGCR), one of the key limiting enzyme involved in cholesterol synthesis (which is also the target of the statin, an anti-cholesterol), is upregulated when *jouvence* is overexpressed. Inversely, it is downregulated when *jouvence* is depleted, suggesting that this crucial gene, involved in the regulation of the cholesterol level, is likely directly regulated by *jouvence*.

Several snoRNAs (both C/D and H/ACA boxes) have also been involved in many biological cancer processes, as tumor initiation, invasion, metastasis, and/or proliferative signalling [10,11]. For examples, some snoRNAs have been associated with the p53 pathway, a well-known tumor suppressor [54]. Here on HCT116 cells, the p53 gene itself is not directly deregulated, but several of its regulators are affected. Another crucial gene involved in cancer and more particularly in ribosome biogenesis is myc [55]. Indeed, in breast cancer, the Myc oncogene upregulates the expression level of fibrillarin, an enzymatic small nucleolar ribonucleoprotein (snoRNP), which led to elevated snoRNAs biogenesis, and consequently inducing p53 suppression [56]. Interestingly, in various model organism, as *C. elegans* [57], *Drosophila* [58], and mouse [59], a link between Myc, the ribosomes biogenesis, and lifespan has been established. Furthermore, in *Drosophila*, it has been proposed that snoRNAs are a novel class of biologically relevant Myc targets [60]. Here, in HCT116 cells overexpressing *jouvence*, MYC and MYCL are upregulated as well as some of its multiple regulators/interactors. Moreover, the depletion of some snoRNAs, as U3 and U8 led to ribosomes dysfunction [61]. Interestingly, the ribosome biogenesis is strongly affected in *jouvence* depleted HCT116 cells. Finally, the SNORD76, a box C/D snoRNA has been shown to act as a tumor suppressor in glioblastoma [62], a similar effect observed with the depletion of *jouvence* in U87-MG glioblastoma cell line. In conclusion, as previously suggested [10,11], snoRNAs have the potential to become cancer biomarkers, and even may potentially become major cancer therapeutic targets in the near future. In such way, the snoRNA-*jouvence* is surely another good and promising candidate.

## Methods

### Cell lines and culture conditions

Cancer cell lines were obtained from the American Type Culture Collection (Rockville, MD, USA) and were cultured according to the supplier’s instructions. Primary Human Umbilical Vein Endothelial Cells (HUVEC) isolated from the vein of the umbilical cord were obtained from Promocell (Germany). HCT116 cells were cultured in McCoy’s 5A medium supplemented with 10% Fetal Bovine Serum and 1% glutamine. hTERT-RPE1 and Caco2 cells were maintained in Dulbecco’s modified Eagle’s medium: nutrient mixture F-12 supplemented with 10% FBS and 1% glutamine. HEK293 and U87-MG cells were cultured in DMEM 1X medium supplemented with 10 % of FBS and 1% glutamine. MCF7 and A549 cells were cultured in RPMI 1640 supplemented with 10% Fetal Bovine Serum and 1% glutamine. HUVECs were grown in Endothelial Cell Growth Medium 2 which is low-serum (2% V/V) media optimized for the cultivation of endothelial cells from large blood vessels. Cells were incubated at 37°C in a humidified atmosphere containing 5% CO2.

### Stable cell transfections

In order to generate stable cell lines that overexpress the human snoRNA-*jouvence*, the *h-jou* was cloned in the pcDNA3.1 Zeo + plasmid (ThermoFisher Scientific, USA) between EcoRI sites, in 5’ - 3’ direction behind a T7 promoter. HCT116, Caco2 or HEK 293 cells were transfected with the empty plasmid, serving as a control or with the plasmid containing *h-jou* cloned sequence. Briefly, cells were plated in 12-well plates and allowed to grow 24 hours and reach nearly 60 to 70% of confluence at the time of transfection. Cells were transfected with the corresponding plasmid, using the Lipofectamine 3000 (ThermoFisher Scientific, USA) at a ratio of 1:2 for the DNA:lipofectamine. Lipofectamine 3000, p3000 and the plasmids were previously diluted in OptiMEM 1X (Gibco) (a Reduced-Serum Medium). The medium was changed 24 hours after the transfection, and cells were allowed to grow for another additional 24 hours before being split to a lower concentration. 72 hours post-transfection, cells were treated with Zeocine (ThermoFisher Scientific) (75 µg/mL, 200 µg/mL, and 150 µg/mL for the HCT116, Caco2 and HEK 293 cells respectively). The selection of stably transfected cells was conducted over a period of nearly 4 to 5 weeks with the addition of Zeocine every 3 to 4 days. Stable clones expressing either the empty plasmid or the plasmid with the human snoRNA were selected, verified by RT-qPCR (TaqMan) before their use in the different experiments.

### Lentivirus preparation and *in-vitro* infection (transduction)

Classical Lentiviral integrative vector expressing the snoRNA-*jouvence* placed downstream to the U6 promoter (pLV.U6.hsnoRNA-jouvence), and containing the puromycine selection marker was generated by Flash Therapeutics/Vectalys (Toulouse, France). Cell infections were carried out according to Flash Therapeutics/Vectalys recommendations. For the HCT116 cell line, two MOI have been tested (MOI-1 and MOI-10). Since MOI-10 gives good results and the highest snoRNA-jou expression level, we uses MOI-10 for all other cell lines. For all cell lines, a puromycine selection has been performed for 48 hours. In such condition, all non-transduced cells were eliminated (performed in parallel on non-lentivirus transduced cells, as control).

### Transfection of siRNA

The effect of the knockdown of the snoRNA-*jouvence* on the HCT116 cells was assessed by the transfection of short interfering RNA (siRNA). First, two different silencer selected siRNAs (Lock-Nucleotid-Acid siRNA or LNA-siRNA, shortly named siRNA) were tested (siRNA-1, antisense sequence: 3’-UCCUCUGUCCACAAUAGCC-5’, Cat nb : 4399665), and (siRNA-2, antisense sequence: 3’-UCAAGACCAAUCACCAUGU-5’, Cat nb : 4399665), as well as a non-targeting siRNA-control used for the specificity of the knockdown (Cat nb: 4390843) (ThermoFisher Scientific, USA). These two siRNAs (as well as the siRNA-control) were independently transfected into the cells in a 12-wells format. Cell suspensions of 0,25 x 10^6^ HCT116 cells per well were directly transfected with the corresponding siRNA (reverse transfection) at a final concentration of 10 nM per well, using the Lipofectamine RNAiMAX transfection reagent (ThermoFisher Scientific, USA). 48 hours after the first siRNA reverse transfection, a forward transfection was performed on the adherent cells (at the same concentration as the first reverse transfection). After 24h, the medium was changed and cells were kept in complete medium. Then, the cells were counted at different time points (days post-infection), depending of the cells lines and compared to the different controls. Cell pellets (for the RNA extraction and RNA-seq analysis) were made 48 hours post-transfection-2 (96h after the first reverse transfection). Conditions were performed in triplicates and the specific knockdown of the snoRNA-*jouvence* was validated by standard RT-qPCR (TaqMan).

### VicellXR counting

The proliferation rate of the stably transfected cells (empty plasmid cells and snoRNA overexpressing cells) was analysed by cell counting. Cells were seeded in 6-well plates (in triplicates), with the following number of cells per well: 0,475 × 10^6^ HCTT16 cells per well; 0,5 × 10^6^ Caco2 cells per well and 0,2 × 10^6^ cells per well for the HEK 293 cells. The experiment was conducted over a period of 96 to 160h. Briefly, the supernatant was harvested and cells washed with PBS 1X. Then, cells were trypsinized with 300 μl of 0.05% trypsin/EDTA (Gibco) per well. Trypsine was inactivated with 700 µL of the corresponding medium per well, and were cells then counted with the VicellXR using trypan blue to determine cell viability (Cell viability Analyzer, Beckman Coulter).

### CellTiter-Glo

A luminescent cell viability assay (CellTiter-Glo, Promega, USA) was assessed to determine the cell proliferation rate based on quantitation of the ATP, an indicator of metabolically active cells. Three thousand cells were plated in each well of a 96-well white plate with clear bottom (Costar 3610, Corning Incorporated, USA). Each cell condition was plated in 6 or 10 wells. Cells were allowed to adhere for 24h, before counting. The homogeneous assay procedure involves addition of 100 µL of the CellTiter-Glo reagent directly to the cells cultured in 100 µL of their corresponding complete medium, before measuring the relative luminescence using a multiplate reader (POLAR Star Omega BMG LABTECH). The experiment was conducted over a week, with counting performed every 24h for HCT116 and HEK293, and at 96 hours for the lentivirus transduction.

### RNA extraction and Reverse Transcription (RT)

Total RNA from human cell lines were extracted using NucleoSpin RNA Plus columns (Macherey-Nagel, France), according to the manufacturer’s instructions, as in Soulé et al., 2020 [13]. Extracted RNAs were verified for the absence of genomic DNA contamination, by performing a RT-PCR using the ribosomal gene RP49. Contaminated samples were therefore treated with RQ1 DNase (Promega, USA) and cleaned with the NucleoSpin RNA Clean-UP (Macherey-Nagel). Two micrograms of total RNA were used for the synthesis of cDNA with oligo-dT (used for Sybr Green RT-qPCR) or random primers (used for TaqMan RT-qPCR for the detection of the snoRNA expression). The M-MULV reverse transcriptase (Promega, USA) was used and the RT was performed in a final volume of 25 µL. The thermal cycling conditions were 37°C for 50 min followed by 15 min at 70°C. RNase H (Invitrogen, USA) was performed to digest RNA-DNA hybrids.

### RT-qPCR (Real Time Quantitative Polymerase Chain Reaction) on selected genes

The differential expression of the snoRNA and selected genes was analysed by real-time PCR (QuantStudio 3, Applied Biosystems, France). The expression of the snoRNA was detected with TaqMan customized probes (Applied Biosciences, Life Technologies) and the TaqMan Universal Master Mix II, no UNG (Applied Biosystems). All conditions were normalized to the GAPDH control gene. The Sybr Green RT-qPCR was performed for all the other genes with the Power UP SYBR-Green PCR Master Mix, according to the manufacturer’s instructions (Applied Biosystems). Primers were designed using the “Primer 3 Plus” software (the primer sequences will be provided upon request). All conditions were normalized to the RPLP0 control gene. The results were analysed using the 2-ΔΔCt method, and displayed as the fold change compared to the control gene.

### Transcriptomic analysis (RNA-seq)

The Transcriptomic analysis (RNA-seq) was performed by Novogene (China). Briefly, according to Novogene, total RNA was extracted and after sample quality control, libraries were generated and checked for quality. Then the libraries were sequenced on an Illumina Hiseq platform and 125 bp/150 bp paired-end reads were generated. The resulting data were controlled, and analyzed with bioinformatic tools. The reference genome and gene model annotation files were downloaded from genome website directly. Index of the reference genome was built using Bowtie v2.2.3 and paired-end clean reads were aligned to the reference genome using TopHat v2.0.12. Novogene uses TopHat as the mapping tool because TopHat can generate a database of splice junctions based on the gene model annotation file and thus a better mapping result than other non-splice mapping tools. For the quantification of gene expression level, Novogene uses HTSeq v0.6.1 to count the reads numbers mapped to each gene. Then FPKM of each gene was calculated based on the length of the gene and reads count mapped to this gene. FPKM, expected number of Fragments Per Kilobase of transcript sequence per Millions base pairs sequenced, considers the effect of sequencing depth and gene length for the reads count at the same time, and is currently the most commonly used method for estimating gene expression levels. Novogene uses DESeq R package (1.18.0) to perform differential expression analysis of two conditions (three biological replicates per condition). DESeq provide statistical routines for determining differential expression in digital gene expression data using a model based on the negative binomial distribution. Prior to differential gene expression analysis, for each sequenced library, the read counts were adjusted by edgeR program package through one scaling normalized factor. The P values were adjusted using the Benjamini & Hochberg method. Corrected P-value of 0,05 and log2 (Fold-change) of 1 were set as the threshold for significantly differential expression. The biological variation was eliminated (case with biological replicates), and the threshold was therefore normally set as p adjusted <0,05. Gene Ontology (GO) enrichment analysis of differentially expressed genes was implemented by the GOseq R package, in which gene length bias was corrected. GO terms with corrected p-value less than 0,05 were considered significantly enriched by differential expressed genes. For the KEGG database [14,15], Novogene uses KOBAS software to test the statistical enrichment of differential expression genes in KEGG pathways. We acknowledge Novogene (China) for the assistance in the Method descriptions of the transcriptomic analysis.

### Statistical Analysis

All data were analysed statistically using one-tailed unpaired t-test, with GraphPad Prism™ software.

## Acknowledgements

We thank K. Sidelarbi and L. Mellottée for their technical assistance. This work was supported by Ninovax (a subsidiary of Truffle Capital, Paris), and by the CNRS to J. Bignon and JR Martin.

## Author contributions

JRM and JB conceived and designed the experiments. FEK and JB performed experiments. FEK, JB, JRM analysed the data. JRM wrote the manuscript with input from all authors.

## Supplementary Information

One Supplementary Figure and eight Supplementary Tables (S1 to S8) containing the RNA-Seq Data-analysis are available (Excel files).

## Supplementary Information

**Suppl. Figure 1).**
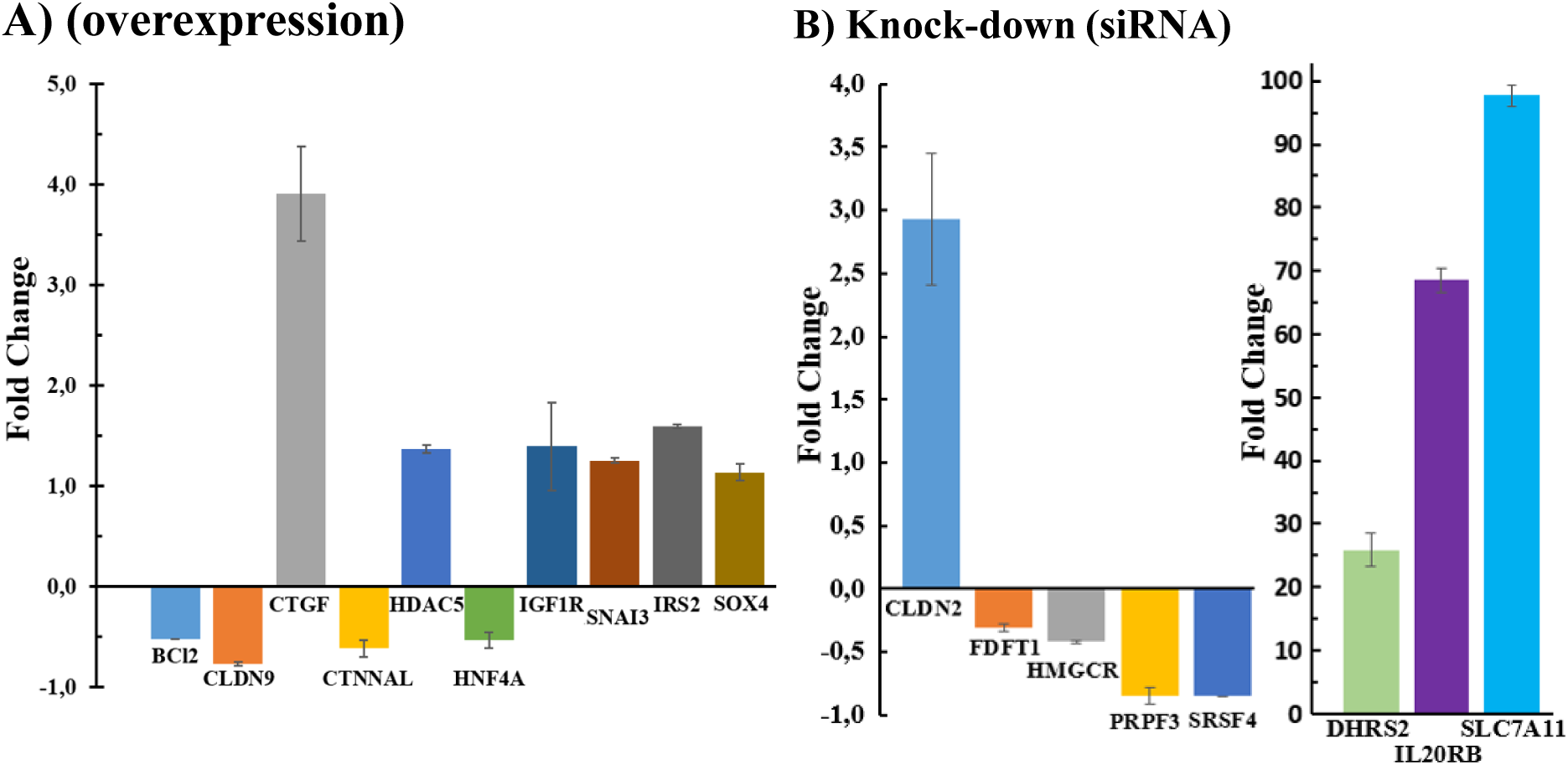
Validation by RT-qPCR of few selected deregulated genes in *jou*overexpression or knockdown. **A)** RT-qPCR (SybGreen) results of the quantification of few selected genes in *jou* overexpressing HCT116 cells. Fold change comparing HCT116 transfected cells versus empty plasmid cells. As revealed by the RNA-Seq, the genes BCL2, CLDN9, CTNNAL, and HNF4A are downregulated, while the genes CTGF, HDAC5, IGF1R, SNAI3, IRS2, and SOX4 are upregulated. **B)** RT-qPCR (SybGreen) results of the quantification of few selected genes in *jou* knockdown. Fold change comparing HCT116 siRNA transfected cells versus non-transfected cells. FDFT1, HMGCR, PRPF, SRSF4 are downregulated, while CLDN2, DHRS2, ILORB, and SLC7A11 genes are upregulated (n=2). The RT-qPCR confirms the deregulation of those genes revealed by RNA-Seq.

### Supplementary Tables S1 to S8

Eight Supplementary Tables (S1 to S8) containing the RNA-Seq Data-analysis (Excel files).

Table S1-HCT-jou-overexpression_DEG upregulated genes.xlsx

Table S2-HCT-jou-overexpression_DEG downregulated genes.xlsx

Table S3-HCT-jou-overexpression_KEGG enrichment pathways.xlsx

Table S4-HCT-siRNA knockdown_DEG upregulated genes.xlsx

Table S5-HCT-siRNA knockdown_DEG downregulated genes.xlsx

Table S6-HCT-siRNA knockdown_KEGG enrichment pathways.xlsx

Table S7-jou-overexpression-genes Up vs siRNA-genes Down-1102 common genes.xlsx

Table S8-jou-overexpression-genes Down vs siRNA-genes Up-868 common genes.xlsx

